# Microbially Produced Bile Acids are Associated with High Levels of IgG Autoantibodies and Worse Mental Well-being in Fibromyalgia Subjects

**DOI:** 10.1101/2024.10.17.618791

**Authors:** Jenny E. Jakobsson, Henrik Carlsson, Ida Erngren, Joana Menezes, Emerson Krock, Matthew A. Hunt, Jeanette Tour Sohlin, Asma Al-Grety, Katalin Sandor, Eva Kosek, Camilla I. Svensson, Kim Kultima

## Abstract

Fibromyalgia (FM) is a disease primarily associated with chronic widespread pain, but other common symptoms are anxiety and depression. We previously proposed that autoimmunity contributes to FM based on findings of increased immunoglobulin G binding to satellite glial cells (anti-SGC IgG) in FM subjects compared to healthy controls (HC). Emerging research suggests that an altered gut microbiota composition is connected to psychological symptoms in FM rather than pain. Gut microbiota can produce or alter bile acids (BAs) and short-chain fatty acids (SCFAs), which have immune and inflammatory functions. Here, we investigate alterations in BA and SCFA concentrations in FM subjects compared to healthy controls (HC) and potential associations with FM symptoms and anti-SGC IgG levels.

Bile acids and SCFAs were quantified using liquid chromatography coupled with high-resolution mass spectrometry and anti-SGC IgG levels were assessed with immunocytochemistry. The correlations between FM symptoms, anti-SGC IgG levels, and serum concentrations of 24 BAs and 11 SCFAs in 35 FM subjects and 32 matched HC were examined.

Fibromyalgia subjects had significantly higher levels of microbially produced BAs than HC. Strikingly, 11 out of 24 BAs were significantly elevated in FM subjects with high, compared to those with low, anti-SGC IgG levels. Concentrations of specific BAs were associated with increased disease severity and worse mental well-being.

These results revealed increased levels of secondary BAs in FM subjects compared to HC. The strong association between BAs, anti-SGC IgG levels, and mental well-being may help elucidate the importance of BAs in the psychological symptoms of FM.

**Significance statement:** Fibromyalgia (FM) is a chronic pain syndrome that significantly impacts an individual’s quality of life. In addition to persistent pain, people with FM often experience depression, anxiety, and irritable bowel syndrome. Previously, we identified autoantibodies that bind to satellite glial cells in the dorsal root ganglia associated with more severe FM symptoms. Our current results demonstrate a novel association between these autoantibodies in FM and bile acids (BAs). Bile acids are essential for lipid metabolism but also act as signaling molecules. We show that patients with elevated levels of autoantibodies also exhibit increased levels of BAs. Furthermore, the BAs strongly correlate with worse FM symptom severity, particularly affecting mental well-being. Our study suggests that lowering BA levels could alleviate the psychological symptoms associated with FM.

## Introduction

Fibromyalgia (FM) ranks among the primary causes of chronic widespread pain (Sarzi-Puttini et al., 2020), with a general prevalence estimated at 2 - 5% and a female preponderance (Branco et al., 2010; Heidari et al., 2017). Fibromyalgia and irritable bowel syndrome (IBS) are examples of frequently co-existing nociplastic pain conditions that are typically associated with symptoms such as disturbed sleep, fatigue, cognitive dysfunction, depression, and anxiety (Fitzcharles et al., 2021; Kosek, 2024, *In press*). While FM has traditionally not been considered an autoimmune disorder, our recent findings suggest that autoimmunity may play a role in the symptomatology of FM. Immunoglobulin G (IgG) antibodies from FM subjects were transferred into mice and induced FM-like symptoms (Goebel et al., 2021). These mice exhibited symptoms such as heightened sensitivity to painful stimuli and reduced intraepidermal nerve fiber density (IENFD), consistent with previous observations in FM patients (Evdokimov et al., 2019; Goebel et al., 2021).

Furthermore, the FM IgG was found to bind to both human and murine satellite glial cells (SGCs) in the dorsal root ganglia (DRGs) (anti-SGC IgG). Stressing the clinical relevance, the anti-SGC IgG levels from FM patients were associated with pain intensity and disease severity (Krock et al., 2023).

Gastrointestinal symptoms are common among FM subjects, with up to 70% experiencing comorbid IBS (Wallace and Hallegua, 2004). Several studies have documented alterations in the gut microbiome composition between FM subjects and healthy controls (HC) (Clos-Garcia et al., 2019; Minerbi et al., 2019, 2023; Kim et al., 2023), along with increased intestinal epithelial permeability (Goebel et al., 2008). Interestingly, fecal microbiota transplantation (FMT) has been shown to alleviate pain and depression symptoms in FM and IBS (El-Salhy et al., 2020; Lin et al., 2021; Cai et al., 2023; Fang et al., 2024). Some overlap with changes in the gut microbiota composition of FM patients, and increased intestinal permeability have been noted in subjects with IBS (Zhou et al., 2009; Garofalo et al., 2023), which may indicate a connection between the etiology of FM and IBS.

Bile acids (BAs) are steroid acids essential for lipid absorption but also participate in inflammatory and immune processes (Ticho et al., 2019; Fuchs and Trauner, 2022). They play significant roles in gastrointestinal disorders like bile acid diarrhea (BAD), which in turn is a common comorbidity in diarrhea-predominant irritable bowel syndrome (IBS-D) (Slattery et al., 2015; Camilleri and Nurko, 2022). Chenodeoxycholic acid (CDCA) and cholic acid (CA), named primary BAs, are synthesized from hepatic cholesterol and converted by gut microbiota into secondary BAs, such as deoxycholic acid (DCA) and lithocholic acid (LCA) (Fuchs and Trauner, 2022). Bile acid synthesis is primarily regulated by a feed-forward loop, where BAs bind to the farnesoid X receptor (FXR) to inhibit further synthesis (Chiang, 2009). The FXR and another BA receptor, takeda G protein-coupled receptor 5 (TGR5), have shown anti-inflammatory effects (Wang et al., 2008; Yoneno et al., 2013), while BAs themselves have been seen to promote inflammation (McMillin et al., 2017; Ju et al., 2023), reflecting the diverse role of BAs in this process.

Bile acids may have central and peripheral functions, as they have been found in cerebrospinal fluid and can cross the blood-brain barrier (Hurley et al., 2022; Harnisch et al., 2023). Minerbi et al. reported that the secondary BA α-muricholic acid (α-MCA) was depleted in overweight individuals with FM and negatively associated with widespread pain index and scores of the fibromyalgia impact questionnaire (FIQ). However, the levels of autoantibodies in FM or questionnaires used in clinical settings to assess depression or anxiety were not included in this study (Minerbi et al., 2023).

The gut microbiota also produces short-chain fatty acids (SCFAs) through fermentation of dietary fibers or protein catabolism. Altered concentrations of SCFAs, specifically butyric acid and propionic acid, have been found in serum from FM subjects compared to HC (Minerbi et al., 2019). However, associations between SCFA concentrations and FM symptoms were not performed. Short-chain fatty acids have been suggested to be important in neuropathic pain and autoimmune conditions, such as inflammatory bowel disease (IBD) (Zhou et al., 2021; Rasouli-Saravani et al., 2023). However, the relationship between the SCFAs and autoimmunity in FM remains unexplored. This study aimed to investigate the associations between anti-SGC IgG levels and BAs and SCFAs concentrations, respectively, in normal-weight FM subjects and healthy controls (HC). Furthermore, we examined how these associations relate to symptoms of FM, assessed with several established questionnaires.

## Methods

### Study participants

The participants of this study were 35 FM subjects and 32 HC, non-overweight (body mass index (BMI) < 25) females aged 20 - 60 years from a larger cohort which has been previously described (Sandström et al., 2020; Ellerbrock et al., 2021a, 2021b; Fanton et al., 2021, 2023; Tour et al., 2022). The local ethical review committee has approved the study (no. 2014/1604-31/1).

The included FM subjects were required to fulfill both the 1990 (Wolfe et al., 1990) and 2011 (Wolfe et al., 2011) American College of Rheumatology (ACR) FM classification criteria, as verified by a pain specialist. All participants signed written consent. Individuals were omitted if any of the following criteria were fulfilled: hypertension (> 160/90 mmHg), pregnancy, autoimmune or pain disorders (apart from FM), neurological or severe somatic disorders, diabetes, psychiatric disorders (such as depression or anxiety), smoking more than five cigarettes per day, substance abuse, current antidepressant or anticonvulsant use, current treatment for anxiety or depression, or not speaking Swedish. Additionally, the patients were required to refrain from analgesics, nonsteroidal anti-inflammatory drugs (NSAIDs), or hypnotics for 48 hours before participation. The HC were enlisted, provided they were free from the criteria above and devoid of chronic pain and confirmed not to take regular sleep medication, antidepressants, NSAIDs, analgesics, or anticonvulsants.

The number of months with pain (pain duration) and FM (FM duration) and total FIQ scores were noted for the FM subjects. Pain intensity ratings using the visual analogue scale (VAS), hospital anxiety and depression (HAD) scales, and short form 36 health survey questionnaire (SF-36) were filled by both FM subjects and HC. The FIQ assesses the disability of FM patients, where a higher score reflects a higher disability (Burckhardt et al., 1991). The pain intensity ratings were assessed with a VAS for weekly average pain (VASavr) and maximum pain during the last week (VASmax). The HAD scales assess depression (HAD D) and anxiety (HAD A), where a higher score indicates a higher likelihood of the condition (Zigmond and Snaith, 1983). The SF-36 has eight categories: general health (GH), physical functioning (PF), physical role functioning (RP), emotional role functioning (RE), social functioning (SF), bodily pain (BP), mental health (MH), and vitality (VT) (Ware and Sherbourne, 1992). The summary of SF-36 GH, PF, RP, and BP was named the physical component of the SF-36 (SF-36 PCS). The SF-36 RE, SF, MH, and VT were summarized to make up the mental component of the SF-36 (SF-36 MCS). A low SF-36 score indicates a higher disability, while the highest indicates no disability.

### Assessment of anti-SGC IgG levels

The anti-SGC IgG levels were estimated through immunocytochemistry. The protocol has been described previously (Krock et al., 2023) and the anti-SGC IgG dataset has been published before (Fanton et al., 2023). Blood samples from FM subjects and HC were collected from the median antecubital vein to a BD vacutainer® STT II tube. The serum was collected by aliquoting the supernatant after blood centrifugation at 2500 rpm for 10 minutes after being kept at room temperature for 40 minutes and then stored at –80°C. The experimenters were blinded to sample belonging during preparation and imaging. A primary cell culture of SGCs was prepared from DRGs harvested from adult female BALB/cAnNRj mice (approved by Stockholm Norra Djurförsöksetiska nämnd). The DRGs were separated by gentle shaking at 37℃ with a papain solution for 30 min, followed by collagenase/dispase solution for an additional 30 min. The cells were resuspended in F12 media with 10% calf serum and 1x penicillin-streptomycin. The cell culture was then triturated, filtered through a 100 μm cell strainer, and supplied to a Nunc™ Lab-Tek™ chamber slides. After 1.5 h, the supernatant was removed to discard non-SGCs. The SGCs were allowed a recovery period overnight in a 5% CO2 incubator at 37℃.

The following day, the frequency of IgG binding to murine SGCs (IgG+SGC%; anti-SGC IgG levels) was assessed. The serum from FM subjects and HC was diluted 1:100 in culture media and filtered with a 0.22 μm filter. After washing, the live SGCs were incubated with the serum IgG for three hours. The SGCs were fixed with 4% formaldehyde for ten minutes, followed by washing with 0.1% Triton-X100 in 1X phosphate buffered saline for 5 minutes. The SGCs were incubated overnight with diluted 1:500 rabbit glutamine synthesis IgG (Abcam ab73593) at 4℃. After the SGC were washed, they were incubated once again but now with anti-human IgG (AF594, Thermo Fisher A11014) and anti-rabbit IgG antibody (AF488, Thermo Fisher A11008) diluted 1:300. Finally, the SGCs were washed, counterstained with Hoescht (10 min), and left to dry (10 min) and coverslipped with Prolong gold mounting media. Imaging was done with a Zeiss LSM800 confocal microscope and then analyzed with a custom machine-learning pipeline (Hunt et al., 2022).

### Mass spectrometry

The concentrations of BAs and SCFAs were determined by high-performance liquid chromatography (HPLC) coupled with high-resolution mass spectrometry (HRMS). Two experimental methods were used, one for BAs and one for SCFAs.

### Sample preparation

#### Bile acids

Fifty µL methanol (MeOH) was added to 25 µL serum from FM subjects and HC. Additionally, 25 µL of a precipitation solution containing isotopically labeled internal standards (IS; all purchased from Avanti polar lipids, Alabaster, AL, USA) was added. The included BA IS were deuterium-labeled Gly-cholic acid (Gly-CA-d4), Gly-ursodeoxycholic acid (Gly-UDCA-d4), Tau-cholic acid (Tau-CA-d4), and Tau-chenodeoxycholic acid (Tau-CDCA-d4). The prepared samples were centrifuged at 21,100 RCF for 15 min at 4°C. The supernatant was then transferred into HPLC vials and stored at –80°C until further analysis.

#### Short-chain fatty acids

Serum samples (20 µL) were precipitated with 80 µL of cold methanol containing IS. The SCFA IS included were deuterium-labeled acetic acid-d3 (Merck, Burlington, MA, United States), propionic acid-d5 (Larodan, Solna, Sweden), butyric acid-d7 (Larodan), and caproic acid-d11 (Larodan). The samples were vortexed and centrifuged at 21,100 RCF for 15 min at 4°C. The supernatant was transferred to a new Eppendorf tube for derivatization. A solution containing 25 mM 3-nitrophenylhydrazine and 25 mM N-(3-Dimethylaminopropyl)-N’-ethylcarboimiide hydrochloride (EDC) was added, followed by a solution of 7% (w/w) of pyridine. The samples were shaken for 30 min at 40°C. The derivatized samples were transferred to HPLC vials and diluted 5.8 times with 0.25% (v/v) formic acid in 1:1 MeOH:water. The samples were stored at −80°C until further analysis.

#### Mass spectrometry acquisition

The samples were analyzed using an Ultimate 3000 HPLC system (Thermo Scientific, Waltham, MA, USA) coupled to a high-resolution hybrid quadrupole Q Exactive Orbitrap MS (Thermo Scientific). All analyses were performed in negative ionization with a resolution of 70,000.

### Bile acids

The samples were injected (20 µL) on a reversed-phase HPLC column (Accucore C18 100 × 2.1 mm, 2.6 µm, Thermo Scientific). A 17.5 min long chromatographic program, including a gradient, was applied using the mobile phases H_2_O with 0.1% acetic acid and 10 mM ammonium acetate (mobile phase A) and MeOH with 0.1% acetic acid and 10 mM ammonium acetate (mobile phase B). The flow rate was 0.6 mL/min, and the column temperature was 55°C. A total of 26 authentic reference standards corresponding to endogenous BAs were used to confirm BA identities: chenodeoxycholic acid (CDCA), cholic acid (CA), deoxycholic acid (DCA), Gly-CDCA, Gly-CA, Gly-ursodeoxycholic acid (Gly-UDCA), hyocholic acid (HCA), hyodeoxycholic acid (HDCA), lithocholic acid (LCA), Gly-LCA 3-sulfate (Gly-LCA 3-S), Tau-CA, Tau-DCA, Tau-LCA, and ursodeoxycholic acid (UDCA) were purchased from Merck; Gly-HCA, Gly-LCA, Tau-HCA, α-muricholic acid (α-MCA), β-MCA, ω-MCA, and murideoxycholic acid (MDCA) were purchased from Cayman Chemical Company (Ann Arbor, MI, United States); Gly-DCA, Gly-HDCA, Tau-CDCA, Tau-HDCA, and Tau-UDCA were purchased from Toronto Research Chemicals (Toronto, ON, Canada). The α-, β- and ω-isomers of MCA could not be chromatographically separated, and the reported MCA is a summarization of these. The HRMS analysis was performed in the full scan mode, collecting data in the range of *m/z* 212.5 – 750 and ionization parameters as described in detail before (Carlsson et al., 2023).

### Short-chain fatty acids

Two µL of sample was injected on a reversed-phase HPLC column (Accucore C18 100 × 2.1 mm, 2.6 µm, Thermo Scientific). A 14-minute long chromatographic program was applied as follows: 2% B for 0.5 min, 2 - 40% B over 5 min, 40 - 100% B over 3 min, 100% B for 2.5 min, and re-equilibration at 2% B for 2 min. The mobile phases contained 0.1% (v/v) formic acid in water (mobile phase A) and 1:9 isopropanol:MeOH (mobile phase B), respectively.

The flow rate was 0.6 mL/min, and the column temperature was 55°C. The HRMS analysis was performed in the full scan mode of *m/z* 100 - 1000 with two selected ion monitoring (SIM) windows: 3.9 - 4.9 min *m/z* 221 - 224 and 4.9 - 6.3 min *m/z* 235 - 238. The spray voltage was 2.4 kV, the capillary temperature and the auxiliary gas heater were set to 320°C and 450°C, the sheath gas flow rate and the auxiliary gas flow rate were set to 55 and 15, respectively, the sweep gas flow rate was 3, and the S-lens RF level was 50.

### Concentration calculations

Calibration curves were generated for SCFAs for which both the native compound and IS were available: 3-hydroxybutyric acid, 2-methylbutyric acid, acetic acid, butyric acid, caproic acid, isobutyric acid, isovaleric acid, propionic acid, and valeric acid. The concentrations of the remaining SCFAs and the BAs were determined through single-point calibration using the IS closest in retention time. The concentrations of acetic acid were adjusted for background levels in the calibrators.

### Statistical analysis

All statistical analyses were performed in R (version 4.3.3). Linear regression (LR) models were used to compare groups with log_2_ transformed concentrations of BAs and SCFAs. All LR models were corrected for age and BMI. First, the models compared FM subjects and HC. Secondly, the models compared FM subjects with high (≥ 50%) IgG+SGC% to low (< 50%) IgG+SGC%.

The BA concentrations were summarized per class: nonconjugated primary, nonconjugated secondary, conjugated (Gly and Tau) primary, and conjugated (Gly and Tau) secondary. The total SCFA concentrations were also summarized. The log_2_ transformed summarized concentrations were compared between FM subjects and HC and FM subjects with high and low IgG+SGC% using a linear model adjusting for age and BMI.

The BA and SCFA concentrations in FM subjects were correlated to the demographic and clinical data collected (age, BMI, pain duration, FM duration, VAS ratings, FIQ, HAD A, HAD D, SF-36 questionnaire, and IgG+SGC%). The correlation analysis for IgG+SGC% was performed with Pearson correlation on log_2_ transformed concentrations. The rest of the clinical data was correlated with Spearman rank’s correlation since they were not normally distributed. P < 0.05 was considered statistically significant.

## Results

### Participant characteristics

Subjects with FM exhibited significantly higher VAS ratings, HAD D scores, and HAD A scores, as well as lower SF-36 PCS and SF-36 MCS scores compared to HC (Table 1). Notably, FM subjects also had elevated levels of anti-SGC IgG (IgG+SGC%). There were no differences in age or BMI between the FM subjects and HC. Pairwise correlation analyses were conducted for parameters in both FM subjects and HC to explore relationships between demographic and clinical data (Figure 1). Anti-SGC IgG levels positively correlated with higher pain intensities in FM subjects. Overall, these findings indicate a poorer quality of life, both physically and psychologically, and increased anti-SGC IgG levels in FM subjects compared to HC.

**Figure 1.**
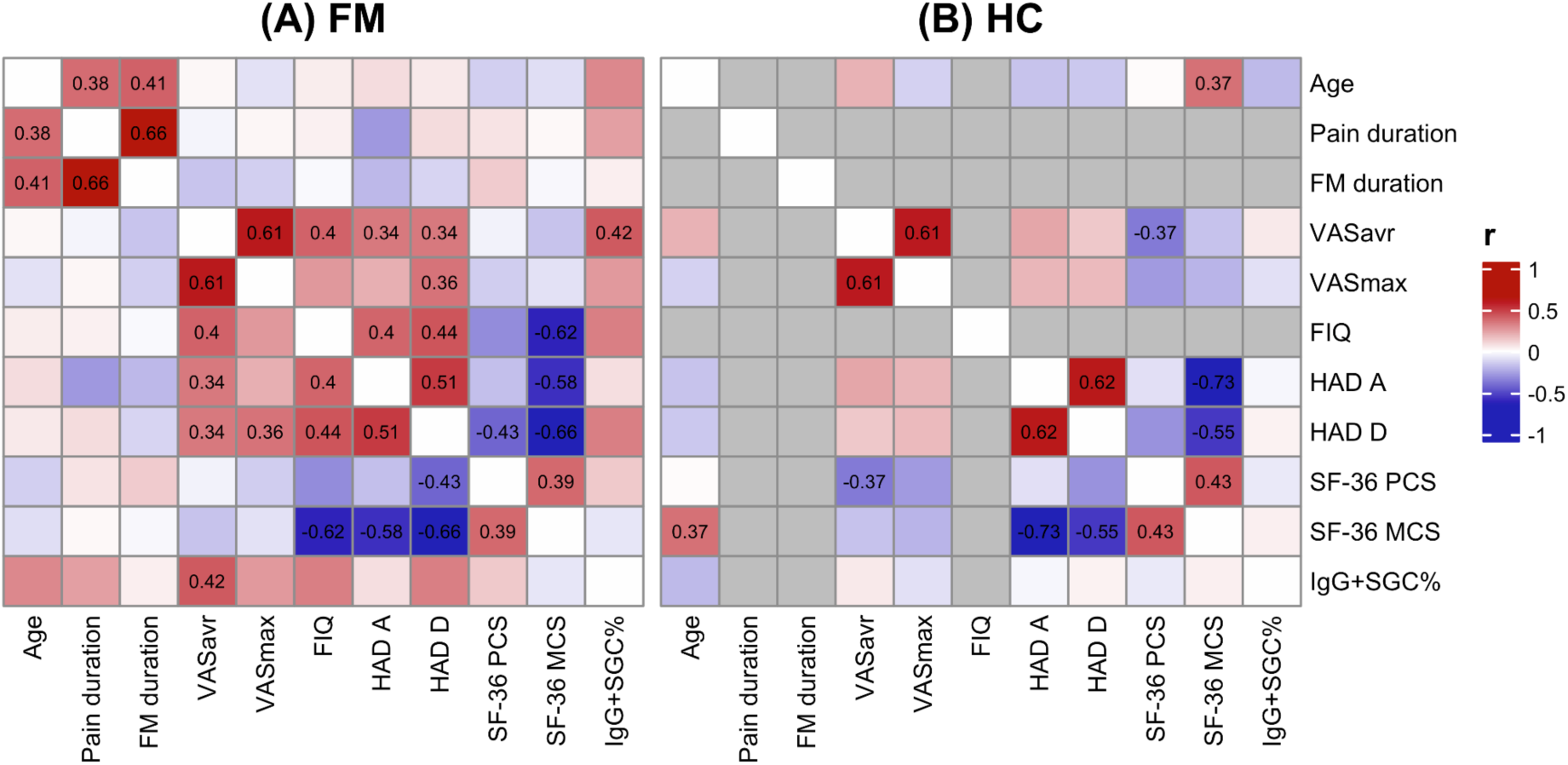
Clinical and demographic data pairwise correlated to each other. The pairwise correlations were done for fibromyalgia (FM) subjects (A) and healthy controls (HC) (B), respectively. A positive correlation is colored in red, and a negative in blue. The numbers shown are the correlation coefficients for significant (P < 0.05) correlations. _VAS: visual analogue scale (pain intensity ratings), VASavr: average weekly pain intensity, VASmax: maximum pain intensity during the past week, FIQ: fibromyalgia impact questionnaire, HAD A/D: Hospital anxiety and depression scale, SF-36: short form 36 health survey questionnaire, SF-36 MCS: mental component of SF-36, SF-36 PCS: physical component of SF-36, IgG+SGC%: frequency of IgG binding to satellite glial cells_

**Table 1.** Descriptive statistics of the fibromyalgia (FM) subjects and healthy controls (HC). The data are presented as mean ± standard deviation. The individuals were divided into HC or FM subjects. The FM subjects were then further divided into low (<50%) or high (≥50%) frequency of immunoglobulin G binding to satellite glial cells (IgG+SGC%). The pain and FM duration (months) and fibromyalgia impact questionnaire (FIQ) scores were only recorded for FM subjects.

|  | HC | FM | Low<br>IgG+SGC% | High<br>IgG+SGC% | FM vs. HC<br>P-value | High vs. low<br>IgG+SGC%<br>P-value |
| --- | --- | --- | --- | --- | --- | --- |
| Samples (n) | 32 | 35 | 19 | 16 | NA | NA |
| Age (year) | 47.8 ± 7.7 | 47.1 ± 8.4 | 45.2 ± 8.5 | 49.4 ± 7.8 | .8456 | .1492 |
| BMI (kg/m<sup>2</sup>) | 22.1 ± 1.5 | 22.5 ± 1.5 | 22.5 ± 1.4 | 22.5 ± 1.7 | .3303 | .8945 |
| Pain duration (months) | NA | 166.8 ± 89.1 | 138 ± 77.5 | 201 ± 92.1 | NA | **.0381\*** |
| FM duration (months) | NA | 117.2 ± 78.5 | 104.7 ± 73.6 | 131.9 ± 83.9 | NA | .6783 |
| VASavr (mm) | 4.9 ± 6.4 | 57.1 ± 18.2 | 50.6 ± 16.9 | 64.9 ± 17 | **<.0001\*** | **.0325\*** |
| VASmax (mm) | 14.3 ± 21.1 | 75.7 ± 14.2 | 72.2 ± 16 | 80 ± 10.7 | **<.0001\*** | .154 |
| FIQ | NA | 61.4 ± 14 | 57.8 ± 12.2 | 65.8 ± 15.2 | NA | .1542 |
| HAD A | 2.9 ± 3 | 7.7 ± 4.7 | 7.6 ± 4.1 | 7.7 ± 5.4 | **<.0001\*** | .702 |
| HAD D | 1.3 ± 1.6 | 6.6 ± 4.2 | 5.5 ± 3.9 | 7.9 ± 4.3 | **<.0001\*** | .07 |
| SF-36 PCS | 93.3 ± 6.5 | 39.6 ± 14.4 | 38.7 ± 9.6 | 40.6 ± 18.9 | **<.0001\*** | .9735 |
| SF-36 MCS | 86.8 ± 12.1 | 52.3 ± 19.9 | 53.9 ± 14.8 | 50.5 ± 25.1 | **<.0001\*** | .9604 |
| IgG+SGC% (%) | 31 ± 18.8 | 48.5 ± 25.6 | 28.3 ± 13.5 | 72.4 ± 11.9 | **.0043\*** | **<.0001\*** |

### Secondary bile acids are elevated in FM subjects compared to HC

When comparing BA concentrations between FM subjects and HC, deoxycholic acid (DCA) was significantly elevated in FM subjects (Table 2). In line with this, the total concentration of nonconjugated secondary BAs was significantly higher in FM subjects than in HC (Table 3). This indicates an altered BA profile in FM subjects, characterized by increased microbially produced secondary BAs.

**Table 2.**
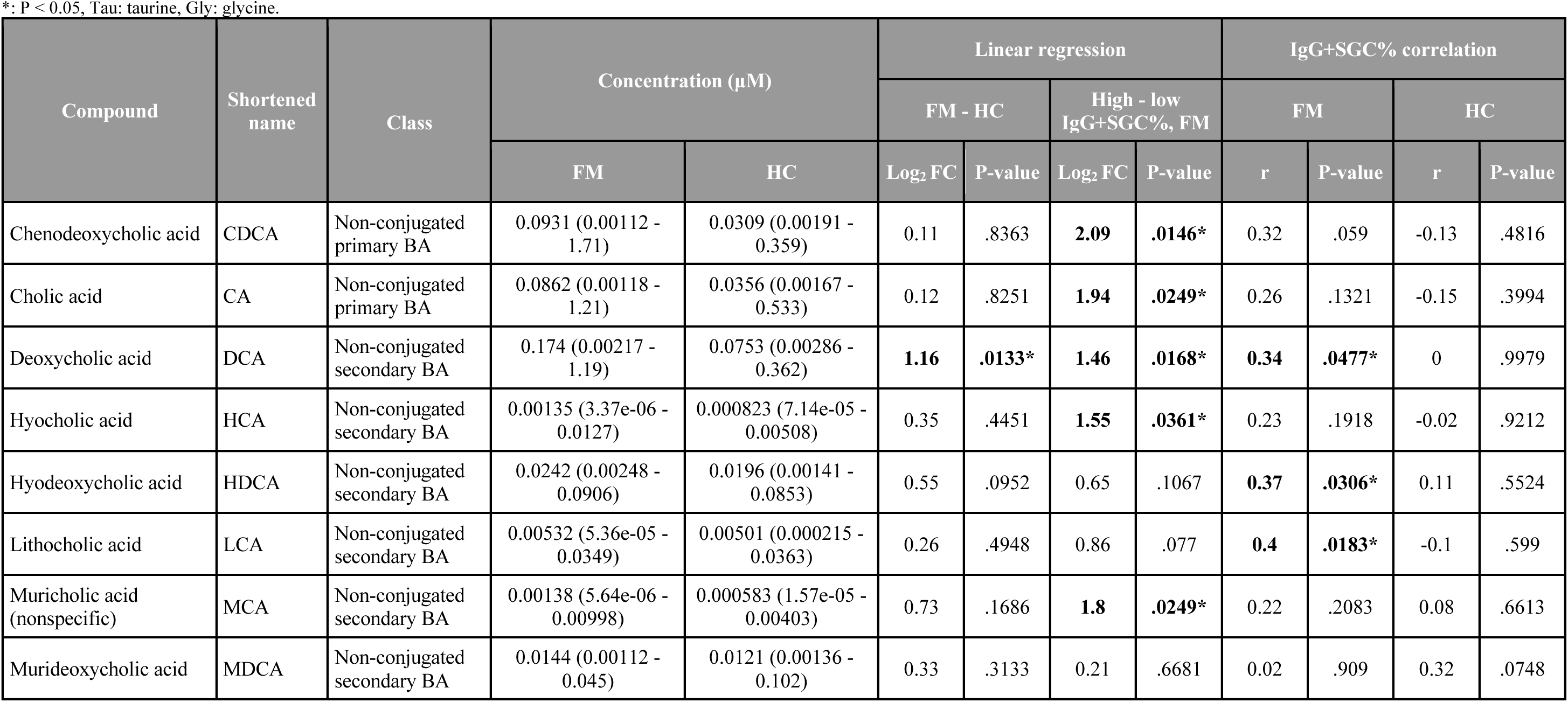
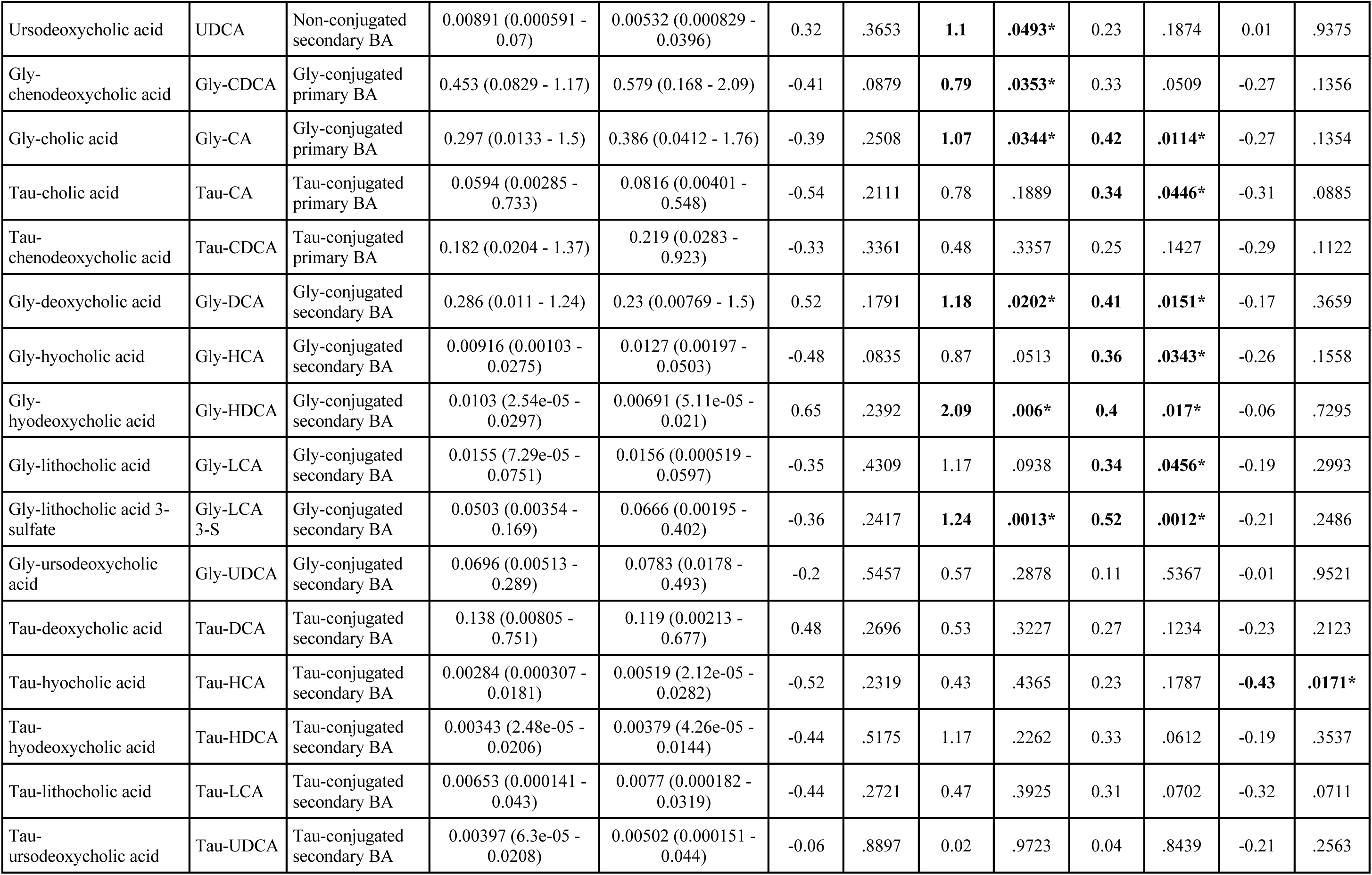
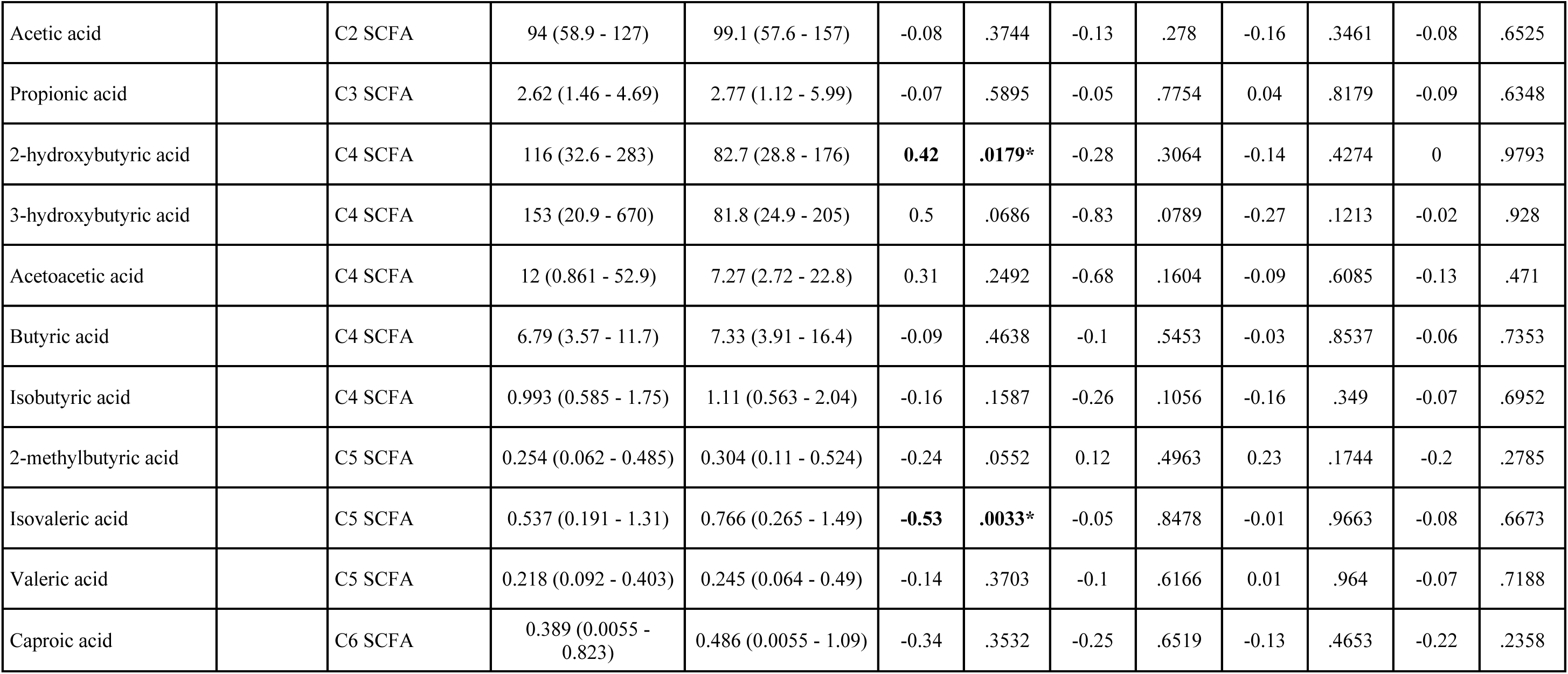
Deoxycholic acid is elevated in fibromyalgia (FM) subjects compared to healthy controls (HC). The concentrations are presented as mean (min - max) in micromolar. The log_2_ transformed fold change (FC) is extracted from the linear regression (LR) model results which was corrected for age and BMI. The correlation analysis of the frequency of IgG binding to satellite glial cells (IgG+SGC%) was performed for FM subjects and HC, respectively.

| Compound | Shortened name | Class | Concentration (μM) |  | Linear regression |  |  |  | IgG+SGC% correlation |  |  |  |
| --- | --- | --- | --- | --- | --- | --- | --- | --- | --- | --- | --- | --- |
|  |  |  |  |  | FM - HC |  | High - low IgG+SGC%, FM |  | FM |  | HC |  |
|  |  |  | FM | HC | Log<sub>2</sub> FC | P-value | Log<sub>2</sub> FC | P-value | r | P-value | r | P-value |
| Chenodeoxycholic acid | CDCA | Non-conjugated primary BA | 0.0931 (0.00112 - 1.71) | 0.0309 (0.00191 - 0.359) | 0.11 | .8363 | **2.09** | **.0146\*** | 0.32 | .059 | -0.13 | .4816 |
| Cholic acid | CA | Non-conjugated primary BA | 0.0862 (0.00118 - 1.21) | 0.0356 (0.00167 - 0.533) | 0.12 | .8251 | **1.94** | **.0249\*** | 0.26 | .1321 | -0.15 | .3994 |
| Deoxycholic acid | DCA | Non-conjugated secondary BA | 0.174 (0.00217 - 1.19) | 0.0753 (0.00286 - 0.362) | **1.16** | **.0133\*** | **1.46** | **.0168\*** | **0.34** | **.0477\*** | 0 | .9979 |
| Hyocholic acid | HCA | Non-conjugated secondary BA | 0.00135 (3.37e-06 - 0.0127) | 0.000823 (7.14e-05 - 0.00508) | 0.35 | .4451 | **1.55** | **.0361\*** | 0.23 | .1918 | -0.02 | .9212 |
| Hyodeoxycholic acid | HDCA | Non-conjugated secondary BA | 0.0242 (0.00248 - 0.0906) | 0.0196 (0.00141 - 0.0853) | 0.55 | .0952 | 0.65 | .1067 | **0.37** | **.0306\*** | 0.11 | .5524 |
| Lithocholic acid | LCA | Non-conjugated secondary BA | 0.00532 (5.36e-05 - 0.0349) | 0.00501 (0.000215 - 0.0363) | 0.26 | .4948 | 0.86 | .077 | **0.4** | **.0183\*** | -0.1 | .599 |
| Muricholic acid (nonspecific) | MCA | Non-conjugated secondary BA | 0.00138 (5.64e-06 - 0.00998) | 0.000583 (1.57e-05 - 0.00403) | 0.73 | .1686 | **1.8** | **.0249\*** | 0.22 | .2083 | 0.08 | .6613 |
| Murideoxycholic acid | MDCA | Non-conjugated secondary BA | 0.0144 (0.00112 - 0.045) | 0.0121 (0.00136 - 0.102) | 0.33 | .3133 | 0.21 | .6681 | 0.02 | .909 | 0.32 | .0748 |
| Ursodeoxycholic acid | UDCA | Non-conjugated secondary BA | 0.00891 (0.000591 - 0.07) | 0.00532 (0.000829 - 0.0396) | 0.32 | .3653 | **1.1** | **.0493\*** | 0.23 | .1874 | 0.01 | .9375 |
| Gly-chenodeoxycholic acid | Gly-CDCA | Gly-conjugated primary BA | 0.453 (0.0829 - 1.17) | 0.579 (0.168 - 2.09) | -0.41 | .0879 | **0.79** | **.0353\*** | 0.33 | .0509 | -0.27 | .1356 |
| Gly-cholic acid | Gly-CA | Gly-conjugated primary BA | 0.297 (0.0133 - 1.5) | 0.386 (0.0412 - 1.76) | -0.39 | .2508 | **1.07** | **.0344\*** | **0.42** | **.0114\*** | -0.27 | .1354 |
| Tau-cholic acid | Tau-CA | Tau-conjugated primary BA | 0.0594 (0.00285 - 0.733) | 0.0816 (0.00401 - 0.548) | -0.54 | .2111 | 0.78 | .1889 | **0.34** | **.0446\*** | -0.31 | .0885 |
| Tau-chenodeoxycholic acid | Tau-CDCA | Tau-conjugated primary BA | 0.182 (0.0204 - 1.37) | 0.219 (0.0283 - 0.923) | -0.33 | .3361 | 0.48 | .3357 | 0.25 | .1427 | -0.29 | .1122 |
| Gly-deoxycholic acid | Gly-DCA | Gly-conjugated secondary BA | 0.286 (0.011 - 1.24) | 0.23 (0.00769 - 1.5) | 0.52 | .1791 | **1.18** | **.0202\*** | **0.41** | **.0151\*** | -0.17 | .3659 |
| Gly-hyocholic acid | Gly-HCA | Gly-conjugated secondary BA | 0.00916 (0.00103 - 0.0275) | 0.0127 (0.00197 - 0.0503) | -0.48 | .0835 | 0.87 | .0513 | **0.36** | **.0343\*** | -0.26 | .1558 |
| Gly-hyodeoxycholic acid | Gly-HDCA | Gly-conjugated secondary BA | 0.0103 (2.54e-05 - 0.0297) | 0.00691 (5.11e-05 - 0.021) | 0.65 | .2392 | **2.09** | **.006\*** | **0.4** | **.017\*** | -0.06 | .7295 |
| Gly-lithocholic acid | Gly-LCA | Gly-conjugated secondary BA | 0.0155 (7.29e-05 - 0.0751) | 0.0156 (0.000519 - 0.0597) | -0.35 | .4309 | 1.17 | .0938 | **0.34** | **.0456\*** | -0.19 | .2993 |
| Gly-lithocholic acid 3-sulfate | Gly-LCA 3-S | Gly-conjugated secondary BA | 0.0503 (0.00354 - 0.169) | 0.0666 (0.00195 - 0.402) | -0.36 | .2417 | **1.24** | **.0013\*** | **0.52** | **.0012\*** | -0.21 | .2486 |
| Gly-ursodeoxycholic acid | Gly-UDCA | Gly-conjugated secondary BA | 0.0696 (0.00513 - 0.289) | 0.0783 (0.0178 - 0.493) | -0.2 | .5457 | 0.57 | .2878 | 0.11 | .5367 | -0.01 | .9521 |
| Tau-deoxycholic acid | Tau-DCA | Tau-conjugated secondary BA | 0.138 (0.00805 - 0.751) | 0.119 (0.00213 - 0.677) | 0.48 | .2696 | 0.53 | .3227 | 0.27 | .1234 | -0.23 | .2123 |
| Tau-hyocholic acid | Tau-HCA | Tau-conjugated secondary BA | 0.00284 (0.000307 - 0.0181) | 0.00519 (2.12e-05 - 0.0282) | -0.52 | .2319 | 0.43 | .4365 | 0.23 | .1787 | **-0.43** | **.0171\*** |
| Tau-hyodeoxycholic acid | Tau-HDCA | Tau-conjugated secondary BA | 0.00343 (2.48e-05 - 0.0206) | 0.00379 (4.26e-05 - 0.0144) | -0.44 | .5175 | 1.17 | .2262 | 0.33 | .0612 | -0.19 | .3537 |
| Tau-lithocholic acid | Tau-LCA | Tau-conjugated secondary BA | 0.00653 (0.000141 - 0.043) | 0.0077 (0.000182 - 0.0319) | -0.44 | .2721 | 0.47 | .3925 | 0.31 | .0702 | -0.32 | .0711 |
| Tau-ursodeoxycholic acid | Tau-UDCA | Tau-conjugated secondary BA | 0.00397 (6.3e-05 - 0.0208) | 0.00502 (0.000151 - 0.044) | -0.06 | .8897 | 0.02 | .9723 | 0.04 | .8439 | -0.21 | .2563 |
| Acetic acid |  | C2 SCFA | 94 (58.9 - 127) | 99.1 (57.6 - 157) | -0.08 | .3744 | -0.13 | .278 | -0.16 | .3461 | -0.08 | .6525 |
| Propionic acid |  | C3 SCFA | 2.62 (1.46 - 4.69) | 2.77 (1.12 - 5.99) | -0.07 | .5895 | -0.05 | .7754 | 0.04 | .8179 | -0.09 | .6348 |
| 2-hydroxybutyric acid |  | C4 SCFA | 116 (32.6 - 283) | 82.7 (28.8 - 176) | **0.42** | **.0179\*** | -0.28 | .3064 | -0.14 | .4274 | 0 | .9793 |
| 3-hydroxybutyric acid |  | C4 SCFA | 153 (20.9 - 670) | 81.8 (24.9 - 205) | 0.5 | .0686 | -0.83 | .0789 | -0.27 | .1213 | -0.02 | .928 |
| Acetoacetic acid |  | C4 SCFA | 12 (0.861 - 52.9) | 7.27 (2.72 - 22.8) | 0.31 | .2492 | -0.68 | .1604 | -0.09 | .6085 | -0.13 | .471 |
| Butyric acid |  | C4 SCFA | 6.79 (3.57 - 11.7) | 7.33 (3.91 - 16.4) | -0.09 | .4638 | -0.1 | .5453 | -0.03 | .8537 | -0.06 | .7353 |
| Isobutyric acid |  | C4 SCFA | 0.993 (0.585 - 1.75) | 1.11 (0.563 - 2.04) | -0.16 | .1587 | -0.26 | .1056 | -0.16 | .349 | -0.07 | .6952 |
| 2-methylbutyric acid |  | C5 SCFA | 0.254 (0.062 - 0.485) | 0.304 (0.11 - 0.524) | -0.24 | .0552 | 0.12 | .4963 | 0.23 | .1744 | -0.2 | .2785 |
| Isovaleric acid |  | C5 SCFA | 0.537 (0.191 - 1.31) | 0.766 (0.265 - 1.49) | **-0.53** | **.0033\*** | -0.05 | .8478 | -0.01 | .9663 | -0.08 | .6673 |
| Valeric acid |  | C5 SCFA | 0.218 (0.092 - 0.403) | 0.245 (0.064 - 0.49) | -0.14 | .3703 | -0.1 | .6166 | 0.01 | .964 | -0.07 | .7188 |
| Caproic acid |  | C6 SCFA | 0.389 (0.0055 - 0.823) | 0.486 (0.0055 - 1.09) | -0.34 | .3532 | -0.25 | .6519 | -0.13 | .4653 | -0.22 | .2358 |

**Table 3.** Fibromyalgia (FM) subjects have increased concentrations of bile acids (BAs) which are associated with the frequency of immunoglobulin G binding to satellite glial cells (IgG+SGC%) compared to healthy controls (HC). The concentrations are presented as mean (min - max) in micromolar. The linear regression results for the summarized groups for short-chain fatty acids (SCFAs) and BAs. The BAs were divided by class.

| Class | Concentration (μM) |  | Linear regression |  |  |  | IgG+SGC% correlation |  |  |  |
| --- | --- | --- | --- | --- | --- | --- | --- | --- | --- | --- |
|  |  |  | FM - HC |  | High - low IgG+SGC%, FM |  | FM |  | HC |  |
|  | FM | HC | Log<sub>2</sub> FC | P-value | Log<sub>2</sub> FC | P-value | r | P-value | r | P-value |
| Total SCFA | 386.594 (144.82 - 1028.08) | 283.886 (166.499 - 544.592) | **0.31** | **.0385\*** | -0.42 | .1098 | -0.24 | .1676 | -0.07 | .7204 |
| Total BA | 1.996 (0.352 - 6.288) | 2 (0.36 - 7.44) | -0.07 | .7937 | **1.13** | **.0025\*** | **0.52** | **.0014\*** | -0.27 | .1369 |
| Non-conjugated primary BA | 0.179 (0.002 - 2.927) | 0.067 (0.005 - 0.893) | 0.14 | .7865 | **2.03** | **.0173\*** | 0.31 | .074 | -0.16 | .3882 |
| Non-conjugated secondary BA | 0.236 (0.028 - 1.312) | 0.122 (0.012 - 0.473) | **0.92** | **.0082\*** | **1.06** | **.0176\*** | **0.41** | **.0147\*** | 0.07 | .7041 |
| Conjugated primary BA | 0.991 (0.137 - 4.693) | 1.265 (0.261 - 4.469) | -0.4 | .1345 | **0.87** | **.0343\*** | **0.4** | **.0176\*** | -0.32 | .0728 |
| Gly-conjugated primary BA | 0.75 (0.096 - 2.593) | 0.965 (0.229 - 3.851) | -0.4 | .1248 | **0.9** | **.0253\*** | **0.4** | **.0164\*** | -0.3 | .094 |
| Tau-conjugated primary BA | 0.241 (0.023 - 2.101) | 0.301 (0.032 - 1.3) | -0.38 | .2831 | 0.56 | .2658 | 0.29 | .0946 | -0.31 | .0888 |
| Conjugated secondary BA | 0.595 (0.063 - 2.364) | 0.549 (0.074 - 2.557) | 0.15 | .6073 | **0.96** | **.0223\*** | **0.42** | **.0125\*** | -0.2 | .275 |
| Gly-conjugated secondary<br>BA | 0.441 (0.038 - 1.545) | 0.409 (0.067 - 2.183) | 0.14 | .6203 | **1.04** | **.0122\*** | **0.43** | **.0094\*** | -0.16 | .3879 |
| Tau-conjugated secondary<br>BA | 0.155 (0.013 - 0.818) | 0.14 (0.006 - 0.73) | 0.32 | .4234 | 0.5 | .335 | 0.28 | .1084 | -0.24 | .1767 |

### Fibromyalgia subjects with high anti-SGC IgG levels have increased bile acid concentrations

The most notable differences in BAs were observed between FM subjects with high *versus* low IgG+SGC% (anti-SGC IgG levels). Fibromyalgia subjects with high anti-SGC IgG levels had significantly higher levels of 11 out of 24 measured BAs compared to those with low anti-SGC IgG levels (Table 2). The BAs significantly elevated were CDCA, CA, DCA, HCA, MCA, UDCA, Gly-CDCA, Gly-CA, Gly-DCA, Gly-HDCA, and Gly-LCA 3-S. Of note, none of the Tau-conjugated BAs showed differences in FM with high IgG+SGC% compared to low.

Summarized by class, FM subjects with high anti-SGC IgG levels had significantly higher total, non-conjugated, and Gly-conjugated BAs, but no difference was seen for Tau-conjugated BAs (Table 3). These findings show that FM subjects with high anti-SGC IgG levels have elevated concentrations of BAs compared to those with low anti-SGC IgG levels.

### Conjugated bile acids are associated with worse mental well-being and increased disease severity in FM subjects

A pairwise correlation analysis was conducted between each clinical parameter and the compound concentrations in FM subjects to investigate potential connections between altered concentrations and FM symptomatology. Conjugated BAs were found to correlate with worse mental well-being. There was a positive correlation between depression symptoms (HAD D) and the concentrations of Gly-DCA, Gly-HDCA, Gly-LCA, Gly-LCA 3-S, Tau-DCA, and Tau-HDCA with correlation coefficients ranging from 0.4 to 0.48 (Supplementary Figure S1). Furthermore, a negative correlation was observed between the mental components of the SF-36 (SF-36 MCS) and the concentrations of Gly-conjugated CA, DCA, HDCA, LCA, and LCA 3-S, as well as Tau-conjugated CA, CDCA, DCA, HCA, HDCA (correlation coefficients ranging from −0.48 to −0.34), exemplified by Gly-DCA in Figure 2. Additionally, Gly-CA, Gly-CDCA, Tau-CA, Tau-CDCA, and Gly-LCA positively correlated with FM disease severity (FIQ), with correlation coefficients ranging from 0.37 to 0.47.

**Figure 2.**
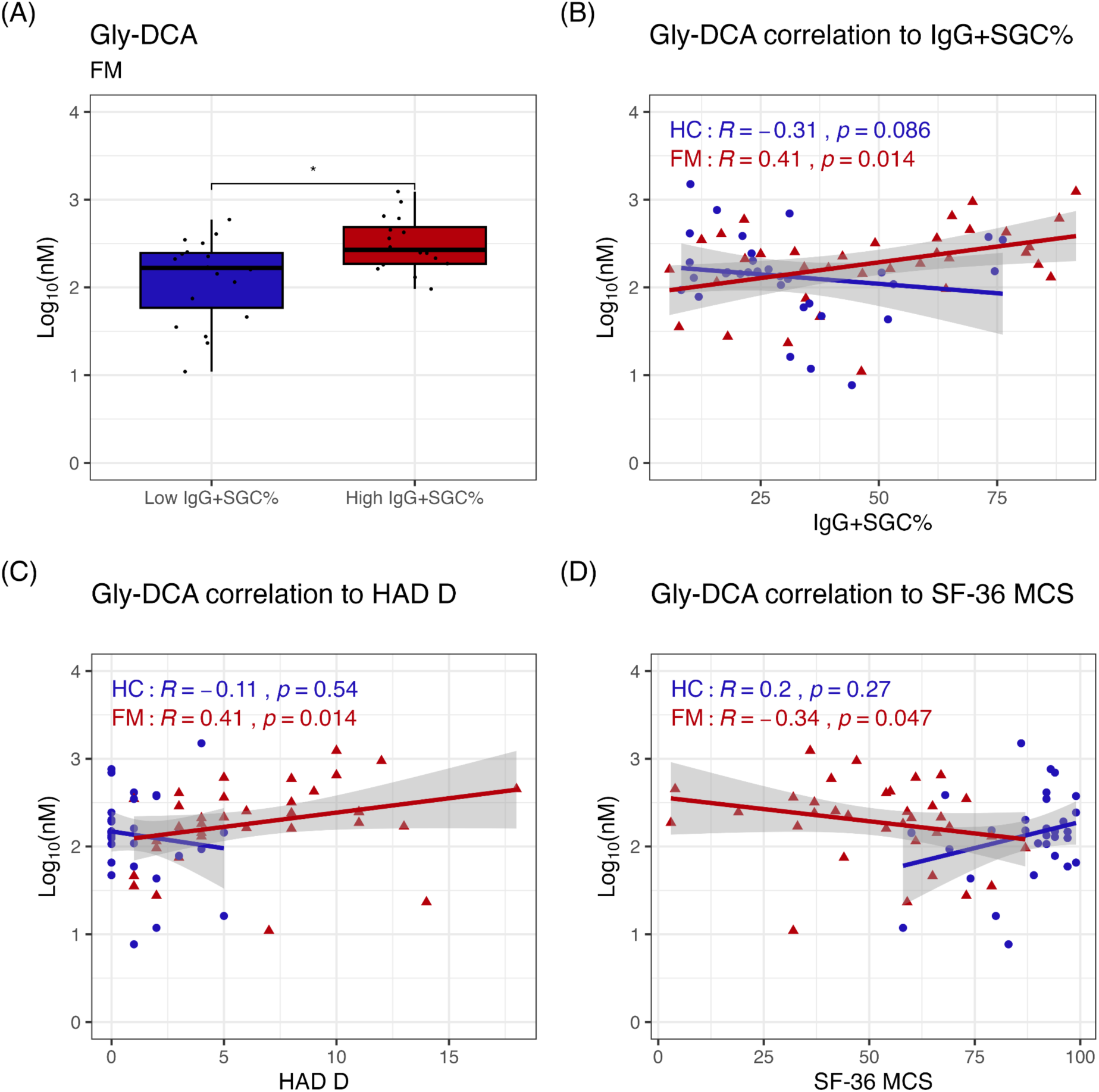
Glycine-conjugated deoxycholic acid (Gly-DCA) is associated with higher anti-SGC IgG levels (IgG+SGC%) and worse mental well-being. Gly-DCA was found in increased concentrations in fibromyalgia (FM) subjects with high (≥50%) IgG+SGC% compared to those with low (<50%) (A). Gly-DCA positively correlated to IgG+SGC% in FM subjects but not in HC (B). Gly-DCA concentrations were associated with depression scores on a Hospital anxiety and depression scale (HAD D) (C) and worse mental well-being (D). _*: P < 0.05, SF-36: short form 36 health survey questionnaire, SF-36 MCS mental summary of SF-36._

Similar results were found when BAs were summarized by class. Conjugated primary and secondary BAs demonstrated negative correlations with SF-36 MCS scores, with correlation coefficients ranging from –0.41 to –0.34 (Figure 3). Total and Gly-conjugated BA concentrations correlated with FIQ scores, with correlation coefficients ranging from 0.34 to 0.44. These findings highlight an association between concentrations of conjugated BAs and worse mental well-being and disease severity in individuals with FM.

**Figure 3.**
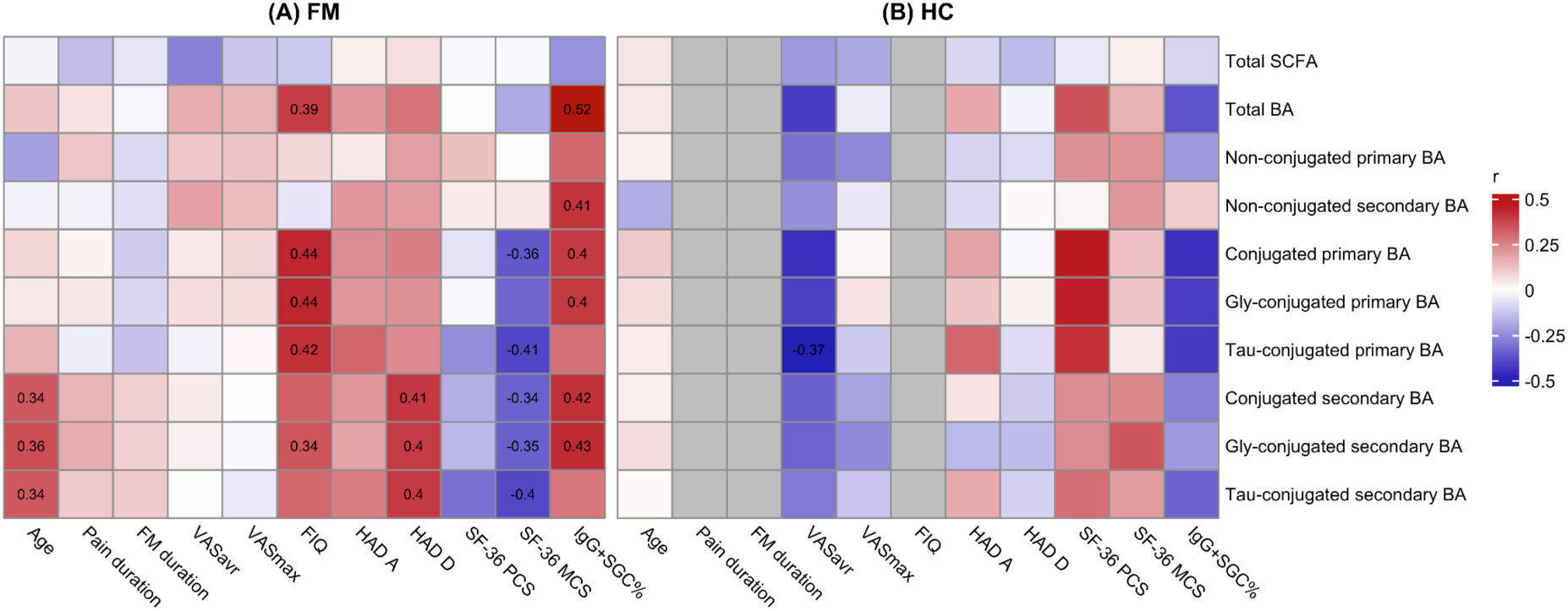
Fibromyalgia (FM) subjects have associations between conjugated bile acids (BAs) and mental well-being. Fibromyalgia subjects’ (A) and healthy controls (HC; B) BA and short-chain fatty acid (SCFA) concentrations were correlated to demographic and clinical data. The frequency of IgG binding to satellite glial cells (IgG+SGC%) was found strongly associated with the total BA, conjugated primary BA, and secondary (non-conjugated and Gly-conjugated) BA concentrations. Conjugated primary and secondary BAs were associated with worse mental well-being (summary of the mental component of the short form 36 health survey questionnaire; SF36 MCS). Conjugated secondary BAs were associated with depression scores on a Hospital anxiety and depression scale (HAD D). A positive correlation is colored in red, and a negative in blue. The numbers shown are the correlation coefficients for significant (P < 0.05) correlation. _Gly: glycine, Tau: taurine, VAS: visual analogue scale (pain intensity ratings), VASavr: average weekly pain intensity, VASmax: maximum pain intensity during the past week, FIQ: fibromyalgia impact questionnaire._

The individual secondary BAs (DCA, HDCA, and LCA) and conjugated BAs (Gly-CA, Tau-CA, Gly-DCA, Gly-HCA, Gly-HDCA, Gly-LCA, and Gly-LCA 3-S) all showed significant positive correlations with anti-SGC IgG levels (correlation coefficient ranging from 0.34 to 0.52) (Supplementary Figure S1). In line with these results, the summarized total, non-conjugated secondary, Gly-conjugated primary, and Gly-conjugated secondary BAs also correlated with anti-SGC IgG levels (correlation coefficient ranging from 0.40 to 0.52) (Figure 3). However, Tau-conjugated BAs did not show these associations. These findings indicate a clear association between increased BA concentrations and anti-SGC IgG levels in individuals with FM.

### Short-chain fatty acid concentrations are altered in FM subjects but not associated with symptom severity

Two SCFAs, isovaleric acid and 2-hydroxybutyric acid, exhibited differences between individuals with FM and HC. We found significantly higher concentrations of 2-hydroxybutyric acid in FM subjects but lower levels of isovaleric acid than HC (Table 2). When summarized, FM subjects showed significantly increased SCFA concentrations compared to HC (Table 3). A significant association between isobutyric acid and VASmax, as well as 2-methylbutyric acid and HAD A, was found in FM subjects. However, we did not find significant correlations between isovaleric acid, 2-hydroxybutyric acid, or total SCFA concentrations with clinical data or FM symptoms (Figure 3, Supplementary Figure S1). In conclusion, FM subjects have altered SCFAs compared to HC, but these changes do not correlate with FM symptoms.

## Discussion

This study explored the intriguing connection between autoimmune mechanisms in FM, changes in BAs and SCFAs, and their associations with FM symptoms. Our findings show elevated levels of secondary BAs in FM subjects compared to HC. Notably, FM subjects with high anti-SGC IgG levels exhibited elevated BA levels compared to those with lower anti-SGC IgG levels. The BA concentrations were strongly associated with poor mental well-being. These findings suggest that BAs contribute to both the psychological symptoms and autoimmune aspects of FM, potentially guiding personalized treatments for different FM subgroups.

### Secondary bile acids in fibromyalgia

Bile acids are humans’ primary product of cholesterol catabolism (Chiang, 2013). There are conflicting reports on total cholesterol levels in FM subjects. Some studies have reported higher total cholesterol levels (Gurer et al., 2006; Loevinger et al., 2007; Cordero et al., 2014), while other studies have found no significant difference between FM subjects and HC (Ozgocmen and Ardicoglu, 2000; Rus et al., 2016). Our analysis shows a notable increase in microbially produced secondary BAs in FM subjects, while the total BA concentrations are unchanged. Rather than an increased cholesterol breakdown, there may be a shift towards secondary BAs. This shift may be attributed to the previously documented gut microbiome alteration in FM subjects (Clos-Garcia et al., 2019; Minerbi et al., 2019; Nhu et al., 2023; Wang et al., 2023). Another possible contributing factor to the increased levels of secondary BAs could be an increased intestinal permeability observed in FM subjects (Goebel et al., 2008; Martín et al., 2023). Taken together, the increased levels of secondary BAs in FM subjects may be due to the reported gut microbiome alteration or an increased influx of these BAs from the intestines.

Previously, α-MCA was reported as decreased in FM subjects compared to HC (Minerbi et al., 2023). In our analysis, we see no significant difference in MCA concentrations. However, our analysis of MCA summarizes the α-, β- and ω-isomers because these isomers could not be chromatographically separated with high confidence, which may be why our results differ. The increased levels of DCA in FM subjects can potentially contribute to inflammation. Low-grade inflammation has been implicated in IBS (Bercik et al., 2005) and FM (Kadetoff and Kosek, 2010). Repetitive instillation of DCA in colon induces low-grade colonic inflammation, leading to persistent visceral hypersensitivity and referred pain in rats (Traub et al., 2008). Thus, there may be a connection between BAs and low-grade inflammation in FM.

### Conjugated bile acids and depression

Fibromyalgia subjects are known to have an increased risk of mood disorders, such as depression (Palomo-López et al., 2019). Our results show a clear association between BAs and psychological symptoms, indicating a connection between BA metabolism and poor mental health in FM subjects. Dysregulation of BAs has been shown in individuals with major depression disorder (MDD), where increased levels of 23-nordeoxycholic acid (norDCA), a DCA derivative, were positively correlated with depression scores (Sun et al., 2022). Interestingly, BA precursors (cholesterol derivatives) have also been found to correlate with depression scores, but not other symptoms, in Parkinsonism (Griffiths et al., 2021).

In both IBD and MDD, alteration of gut microbiome composition and altered secondary BA metabolism have been associated with anxiety and depression scores (Feng et al., 2022; MahmoudianDehkordi et al., 2022). Additionally, a recent study by Nhu and colleagues further highlighted the role of gut-brain axis disruption, possibly tied to BA metabolism in psychological symptoms associated with FM, rather than pain (Nhu et al., 2023). Our results extend this understanding by showing that conjugated primary and secondary BAs are associated with mental health outcomes. However, since hepatic primary BAs show the same associations with mental health outcomes as the microbially produced secondary BAs, changes in the gut microbiome alone may not fully explain the connection. Instead, the alterations in BA metabolism may directly influence depressive symptoms in FM as well as in other diseases like MDD and IBD.

A potential contributor to poor mental well-being in FM subjects may be increased FXR signaling. The farnesoid X receptor has a known affinity for both primary and secondary BAs, including conjugated BAs, which may influence various physiological pathways linked to mental health (Fiorucci et al., 2018; Fuchs and Trauner, 2022). A study in rats showed that stress increased FXR expression in the brain, leading to depressive-like symptoms (Chen et al., 2018). Moreover, FXR knockout mice exhibited increased levels of BAs in serum and brain tissue, along with disruption in neurotransmitter pathways in different brain regions (Huang et al., 2015). In FM patients, increased expression of FXR pathways has been reported (Ramírez-Tejero et al., 2018), with the authors suggesting that this increased activation of FXR pathways may be a response to chronic inflammation. However, as noted by Chen and colleagues (Chen et al., 2018), this increase in FXR signaling could also be connected to depressive symptoms in FM. Further research into the relationship between FXR and BA signaling could shed light on the mechanisms underlying depressive symptoms in FM.

Another BA receptor, TGR5, has also been associated with anti-depressive symptoms, as TGR5 knockout mice exhibit anxiety and depression-like behaviors (Tao et al., 2024). These behaviors were further linked to the gut microbiota, as fecal transplants from TGR5 knockout mice into healthy mice induced similar anxiety- and depression-like symptoms (Tao et al., 2024). Taken together, these findings suggest that BAs and their receptors play a significant role in mental health, with potential implications for both MDD and FM.

### Autoimmunity and bile acids

Previous studies have shown alterations in BA concentrations in autoimmune diseases, such as type 1 diabetes, multiple sclerosis, IBD, and autoimmune hepatitis (Schmucker et al., 1990; Sheng et al., 2017; Bhargava et al., 2020; Feng et al., 2022; Lamichhane et al., 2022). This is particularly interesting, as our findings reveal a strong association between BA levels and the presence of autoantibodies binding to DRG SGCs in FM.

Bile acids can bind to TGR5 on DRG neurons and induce analgesic effects (Alemi et al., 2013). However, BAs can induce visceral hypersensitivity through FXR activation, increasing the expression of transient receptor potential vanilloid 1 (TRPV1) in DRGs (Li et al., 2019). Deoxycholic acid can directly and indirectly stimulate neurons in the colon and DRGs (Yu et al., 2019). In a mouse model of peripheral neuropathic pain, DCA increased neuronal hyperexcitability dependent on TGR5 in DRGs (Zhong et al., 2023). These findings highlight the dual anti- and pro-nociceptive roles of BAs, likely regulated via their receptor interactions. Our results show that DCA and ten other BAs are present at higher concentrations in FM subjects with high anti-SGC IgG levels compared to those with lower levels. These striking findings, along with the close proximity of autoantibody binding and BA activity, suggest a potential overlap or direct interactions between the two. The strong association between anti-SGC IgG and BAs in our study suggests that BAs may contribute to autoimmune mechanisms in FM.

When we further investigated conjugated BAs and their associations with anti-SGC IgG levels, we found that the association depended on the specific amino acid used for conjugation. Glycine-conjugated primary and secondary BAs significantly correlated with anti-SGC IgG levels, whereas the Tau-conjugated BAs showed no such correlations. In humans, Gly-conjugation is the most common BA conjugation, occurring at a 3:1 ratio compared to Tau-conjugation (Warren et al., 2006; Chiang and Ferrell, 2022). Animal studies have shown differences in affinity to TGR5 based on whether BAs are conjugated with glycine, taurine or are non-conjugated, with Tau-conjugated BAs acting as the most potent agonist (Kawamata et al., 2003). Although the role of BA conjugation in chronic pain is not yet defined, our observed association between Gly-conjugated BAs and mental well-being may be due to specific properties of these two different amino acids. Our findings show a novel association between BAs and anti-SGC IgG levels in FM, with conjugation-specific differences that may have important implications for the understanding of the disease.

### Short-chain fatty acid metabolism in fibromyalgia

Isovaleric acid is produced from the branched-chain amino acid leucine through microbial actions (Mortensen and Clausen, 1996). The SCFA-synthesizing bacterial species *Bifidobacterium* has been reported to correlate with isovaleric acid levels in fecal samples of young children (Hemalatha et al., 2017), and its abundance is reduced in FM subjects compared to controls (Clos-Garcia et al., 2019). Interestingly, FM subjects also have increased serum concentrations of leucine compared to HC (Ruggiero et al., 2017; Rus et al., 2024). Unexpectedly, our findings reveal that FM subjects have lower concentrations of isovaleric acid than HC. This discrepancy suggests that the reduced levels of isovaleric acid in FM may be driven by changes in the gut microbiota rather than a decreased availability of leucine.

When glutathione is synthesized in hepatic tissue, 2-hydroxybutyric acid is produced as a byproduct. Notably, the rate of glutathione synthesis increases during oxidative stress (Whiley et al., 2024). The elevated levels of 2-hydroxybutyric acid observed in FM subjects compared to HC may be due to heightened oxidative stress, which has been previously documented in FM (Cordero et al., 2010; Sánchez-Domínguez et al., 2015; Menzies et al., 2020; Dos Santos et al., 2022). To further explore potential disease-related mechanisms, additional research into serum concentrations of SCFA in FM is warranted, particularly since no significant associations with FM symptomatology were identified.

### Limitations

There are several limitations to the conclusions drawn from this study. First, these findings indicate only associations and do not establish casual relationships. Additionally, the study participants were not fasting prior to serum collection, and various lifestyle factors were not recorded, which could potentially influence the serum concentrations of SCFA and BA. Lastly, the study did not assess comorbid IBS, which may also impact the results.

## Conclusion

Our results reveal a strong correlation between BAs and anti-SGC IgG levels in individuals with FM. Notably, conjugated BAs are also associated with poorer mental well-being. This suggests that a disturbed BA circulation in FM may negatively affect mental health and contribute to depressive symptoms, highlighting an essential aspect of the condition that may be linked to the gut-brain axis. Furthermore, bile acids may contribute to both the psychological symptoms and autoimmune aspects of FM, offering potential pathways toward personalized treatment approaches tailored to specific subgroups of FM patients.

## Acknowledgments

Data is available upon reasonable request. This work was supported by funding agents: the European Research Council under the European Union’s Horizon 2020 research and innovation programme grant number 866075; the Knut and Alice Wallenberg Foundation; Swedish Research Council grant numbers 2021-02189 and 2022-00564; a generous donation from Leif Lundblad and family; Region Uppsala (ALF-grant and R&D funds); Region Stockholm (ALF); Swedish Rheumatism Association; Åke Wiberg foundation; FOREUM; FORMAS grant numbers 2020–01267, 2022-00488, and 2023-00905.

## References

Alemi F, Kwon E, Poole DP, Lieu T, Lyo V, Cattaruzza F, Cevikbas F, Steinhoff M, Nassini R, Materazzi S, Guerrero-Alba R, Valdez-Morales E, Cottrell GS, Schoonjans K, Geppetti P, Vanner SJ, Bunnett NW, Corvera CU (2013) The TGR5 receptor mediates bile acid-induced itch and analgesia. J Clin Invest 123:1513–1530.

Bercik P, Verdu EF, Collins SM (2005) Is irritable bowel syndrome a low-grade inflammatory bowel disease? Gastroenterol Clin North Am 34:235–245, vi–vii.

Bhargava P et al. (2020) Bile acid metabolism is altered in multiple sclerosis and supplementation ameliorates neuroinflammation. J Clin Invest 130:3467–3482.

Branco JC, Bannwarth B, Failde I, Abello Carbonell J, Blotman F, Spaeth M, Saraiva F, Nacci F, Thomas E, Caubère J-P, Le Lay K, Taieb C, Matucci-Cerinic M (2010) Prevalence of fibromyalgia: a survey in five European countries. Semin Arthritis Rheum 39:448–453.

Burckhardt CS, Clark SR, Bennett RM (1991) The fibromyalgia impact questionnaire: development and validation. J Rheumatol 18:728–733.

Cai W et al. (2023) Gut microbiota promotes pain in fibromyalgia. bioRxiv:2023.10.24.563794 Available at: https://www.biorxiv.org/content/10.1101/2023.10.24.563794v1.full [Accessed January 12, 2024].

Camilleri M, Nurko S (2022) Bile acid diarrhea in adults and adolescents. Neurogastroenterol Motil 34:e14287.

Carlsson H, Sreenivasan AP, Erngren I, Larsson A, Kultima K (2023) Combining the targeted and untargeted screening of environmental contaminants reveals associations between PFAS exposure and vitamin D metabolism in human plasma. Environ Sci Process Impacts 25:1116–1130.

Chen W-G, Zheng J-X, Xu X, Hu Y-M, Ma Y-M (2018) Hippocampal FXR plays a role in the pathogenesis of depression: A preliminary study based on lentiviral gene modulation. Psychiatry Res 264:374–379.

Chiang JYL (2009) Bile acids: regulation of synthesis. J Lipid Res 50:1955–1966.

Chiang JYL (2013) Bile acid metabolism and signaling. Compr Physiol 3:1191–1212.

Chiang JYL, Ferrell JM (2022) Discovery of farnesoid X receptor and its role in bile acid metabolism. Mol Cell Endocrinol 548:111618.

Clos-Garcia M et al. (2019) Gut microbiome and serum metabolome analyses identify molecular biomarkers and altered glutamate metabolism in fibromyalgia. EBioMedicine 46:499–511.

Cordero MD, Alcocer-Gómez E, Cano-García FJ, Sánchez-Domínguez B, Fernández-Riejo P, Moreno Fernández AM, Fernández-Rodríguez A, De Miguel M (2014) Clinical symptoms in fibromyalgia are associated to overweight and lipid profile. Rheumatol Int 34:419–422.

Cordero MD, De Miguel M, Moreno Fernández AM, Carmona López IM, Garrido Maraver J, Cotán D, Gómez Izquierdo L, Bonal P, Campa F, Bullon P, Navas P, Sánchez Alcázar JA (2010) Mitochondrial dysfunction and mitophagy activation in blood mononuclear cells of fibromyalgia patients: implications in the pathogenesis of the disease. Arthritis Res Ther 12:R17.

Dos Santos JM, Rodrigues Lacerda AC, Ribeiro VGC, Scheidt Figueiredo PH, Fonseca SF, da Silva Lage VK, Costa HS, Pereira Lima V, Sañudo B, Bernardo-Filho M, da Cunha de Sá Caputo D, Mendonça VA, Taiar R (2022) Oxidative Stress Biomarkers and Quality of Life Are Contributing Factors of Muscle Pain and Lean Body Mass in Patients with Fibromyalgia. Biology 11 Available at: 10.3390/biology11060935.

Ellerbrock I, Sandström A, Tour J, Fanton S, Kadetoff D, Schalling M, Jensen KB, Sitnikov R, Kosek E (2021a) Serotonergic gene-to-gene interaction is associated with mood and GABA concentrations but not with pain-related cerebral processing in fibromyalgia subjects and healthy controls. Mol Brain 14:81.

Ellerbrock I, Sandström A, Tour J, Kadetoff D, Schalling M, Jensen KB, Kosek E (2021b) Polymorphisms of the μ-opioid receptor gene influence cerebral pain processing in fibromyalgia. Eur J Pain 25:398–414.

El-Salhy M, Hatlebakk JG, Gilja OH, Bråthen Kristoffersen A, Hausken T (2020) Efficacy of faecal microbiota transplantation for patients with irritable bowel syndrome in a randomised, double-blind, placebo-controlled study. Gut 69:859–867.

Evdokimov D, Frank J, Klitsch A, Unterecker S, Warrings B, Serra J, Papagianni A, Saffer N, Meyer Zu, Altenschildesche C, Kampik D, Malik RA, Sommer C, Üçeyler N (2019) Reduction of skin innervation is associated with a severe fibromyalgia phenotype. Ann Neurol 86:504–516.

Fang H, Hou Q, Zhang W, Su Z, Zhang J, Li J, Lin J, Wang Z, Yu X, Yang Y, Wang Q, Li X, Li Y, Hu L, Li S, Wang X, Liao L (2024) Fecal Microbiota transplantation improves clinical symptoms of fibromyalgia: An open-label, randomized, nonplacebo-controlled study. J Pain 25:104535.

Fanton S, Menezes J, Krock E, Sandström A, Tour J, Sandor K, Jurczak A, Hunt M, Baharpoor A, Kadetoff D, Jensen KB, Fransson P, Ellerbrock I, Sitnikov R, Svensson CI, Kosek E (2023) Anti-satellite glia cell IgG antibodies in fibromyalgia patients are related to symptom severity and to metabolite concentrations in thalamus and rostral anterior cingulate cortex. Brain Behav Immun 114:371–382.

Fanton S, Sandström A, Tour J, Kadetoff D, Schalling M, Jensen KB, Sitnikov R, Ellerbrock I, Kosek E (2021) The translocator protein gene is associated with endogenous pain modulation and the balance between glutamate and GABA in fibromyalgia and healthy subjects: a multimodal neuroimaging study. Pain Available at: 10.1097/j.pain.0000000000002309.

Feng L, Zhou N, Li Z, Fu D, Guo Y, Gao X, Liu X (2022) Co-occurrence of gut microbiota dysbiosis and bile acid metabolism alteration is associated with psychological disorders in Crohn’s disease. FASEB J 36:e22100.

Fiorucci S, Biagioli M, Zampella A, Distrutti E (2018) Bile acids activated receptors regulate innate immunity. Front Immunol 9:1853.

Fitzcharles M-A, Cohen SP, Clauw DJ, Littlejohn G, Usui C, Häuser W (2021) Nociplastic pain: towards an understanding of prevalent pain conditions. Lancet 397:2098–2110.

Fuchs CD, Trauner M (2022) Role of bile acids and their receptors in gastrointestinal and hepatic pathophysiology. Nat Rev Gastroenterol Hepatol 19:432–450.

Garofalo C, Cristiani CM, Ilari S, Passacatini LC, Malafoglia V, Viglietto G, Maiuolo J, Oppedisano F, Palma E, Tomino C, Raffaeli W, Mollace V, Muscoli C (2023) Fibromyalgia and Irritable Bowel Syndrome Interaction: A Possible Role for Gut Microbiota and Gut-Brain Axis. Biomedicines 11 Available at: 10.3390/biomedicines11061701.

Goebel A et al. (2021) Passive transfer of fibromyalgia symptoms from patients to mice. J Clin Invest 131 Available at: 10.1172/JCI144201.

Goebel A, Buhner S, Schedel R, Lochs H, Sprotte G (2008) Altered intestinal permeability in patients with primary fibromyalgia and in patients with complex regional pain syndrome. Rheumatology 47:1223–1227.

Griffiths WJ, Abdel-Khalik J, Moore SF, Wijeyekoon RS, Crick PJ, Yutuc E, Farrell K, Breen DP, Williams-Gray CH, Theofilopoulos S, Arenas E, Trupp M, Barker RA, Wang Y (2021) The cerebrospinal fluid profile of cholesterol metabolites in Parkinson’s disease and their association with disease state and clinical features. Front Aging Neurosci 13:685594.

Gurer G, Sendur OF, Ay C (2006) Serum lipid profile in fibromyalgia women. Clin Rheumatol 25:300–303.

Harnisch L-O, Neugebauer S, Mihaylov D, Eidizadeh A, Zechmeister B, Maier I, Moerer O (2023) Quantification of Bile Acids in Cerebrospinal Fluid: Results of an Observational Trial. Biomedicines 11 Available at: 10.3390/biomedicines11112947.

Heidari F, Afshari M, Moosazadeh M (2017) Prevalence of fibromyalgia in general population and patients, a systematic review and meta-analysis. Rheumatol Int 37:1527–1539.

Hemalatha R, Ouwehand AC, Saarinen MT, Prasad UV, Swetha K, Bhaskar V (2017) Effect of probiotic supplementation on total lactobacilli, bifidobacteria and short chain fatty acids in 2-5-year-old children. Microb Ecol Health Dis 28:1298340.

Huang F, Wang T, Lan Y, Yang L, Pan W, Zhu Y, Lv B, Wei Y, Shi H, Wu H, Zhang B, Wang J, Duan X, Hu Z, Wu X (2015) Deletion of mouse FXR gene disturbs multiple neurotransmitter systems and alters neurobehavior. Front Behav Neurosci 9:70.

Hunt MA, Lund H, Delay L, Dos Santos GG, Pham A, Kurtovic Z, Telang A, Lee A, Parvathaneni A, Kussick E, Corr M, Yaksh TL (2022) DRGquant: A new modular AI-based pipeline for 3D analysis of the DRG. J Neurosci Methods 371:109497.

Hurley MJ, Bates R, Macnaughtan J, Schapira AHV (2022) Bile acids and neurological disease. Pharmacol Ther 240:108311.

Ju J, Zhang C, Yang J, Yang Q, Yin P, Sun X (2023) Deoxycholic acid exacerbates intestinal inflammation by modulating interleukin-1β expression and tuft cell proportion in dextran sulfate sodium-induced murine colitis. PeerJ 11:e14842.

Kadetoff D, Kosek E (2010) Evidence of reduced sympatho-adrenal and hypothalamic-pituitary activity during static muscular work in patients with fibromyalgia. J Rehabil Med 42:765–772.

Kawamata Y, Fujii R, Hosoya M, Harada M, Yoshida H, Miwa M, Fukusumi S, Habata Y, Itoh T, Shintani Y, Hinuma S, Fujisawa Y, Fujino M (2003) A G protein-coupled receptor responsive to bile acids. J Biol Chem 278:9435–9440.

Kim Y, Kim G-T, Kang J (2023) Microbial Composition and Stool Short Chain Fatty Acid Levels in Fibromyalgia. Int J Environ Res Public Health 20 Available at: 10.3390/ijerph20043183.

Kosek E (2024) The concept of nociplastic pain - where to from here? Pain.

Krock E, Morado-Urbina CE, Menezes J, Hunt MA, Sandström A, Kadetoff D, Tour J, Verma V, Kultima K, Haglund L, Meloto CB, Diatchenko L, Kosek E, Svensson CI (2023) Fibromyalgia patients with elevated levels of anti-satellite glia cell immunoglobulin G antibodies present with more severe symptoms. Pain 164:1828– 1840.

Lamichhane S, Sen P, Dickens AM, Alves MA, Härkönen T, Honkanen J, Vatanen T, Xavier RJ, Hyötyläinen T, Knip M, Orešič M (2022) Dysregulation of secondary bile acid metabolism precedes islet autoimmunity and type 1 diabetes. Cell Rep Med 3:100762.

Li W-T, Luo Q-Q, Wang B, Chen X, Yan X-J, Qiu H-Y, Chen S-L (2019) Bile acids induce visceral hypersensitivity via mucosal mast cell-to-nociceptor signaling that involves the farnesoid X receptor/nerve growth factor/transient receptor potential vanilloid 1 axis. FASEB J 33:2435–2450.

Lin H, Guo Q, Wen Z, Tan S, Chen J, Lin L, Chen P, He J, Wen J, Chen Y (2021) The multiple effects of fecal microbiota transplantation on diarrhea-predominant irritable bowel syndrome (IBS-D) patients with anxiety and depression behaviors. Microb Cell Fact 20:233.

Loevinger BL, Muller D, Alonso C, Coe CL (2007) Metabolic syndrome in women with chronic pain. Metabolism 56:87–93.

MahmoudianDehkordi S, Bhattacharyya S, Brydges CR, Jia W, Fiehn O, Rush AJ, Dunlop BW, Kaddurah-Daouk R (2022) Gut Microbiome-Linked Metabolites in the Pathobiology of Major Depression With or Without Anxiety-A Role for Bile Acids. Front Neurosci 16:937906.

Martín F, Blanco-Suárez M, Zambrano P, Cáceres O, Almirall M, Alegre-Martín J, Lobo B, González-Castro AM, Santos J, Domingo JC, Jurek J, Castro-Marrero J (2023) Increased gut permeability and bacterial translocation are associated with fibromyalgia and myalgic encephalomyelitis/chronic fatigue syndrome: implications for disease-related biomarker discovery. Front Immunol 14:1253121.

McMillin M, Frampton G, Grant S, Khan S, Diocares J, Petrescu A, Wyatt A, Kain J, Jefferson B, DeMorrow S (2017) Bile acid-mediated sphingosine-1-phosphate receptor 2 signaling promotes neuroinflammation during hepatic encephalopathy in mice. Front Cell Neurosci 11:191.

Menzies V, Starkweather A, Yao Y, Thacker LR 2nd, Garrett TJ, Swift-Scanlan T, Kelly DL, Patel P, Lyon DE (2020) Metabolomic Differentials in Women With and Without Fibromyalgia. Clin Transl Sci 13:67–77.

Minerbi A, Gonzalez E, Brereton N, Fitzcharles M-A, Chevalier S, Shir Y (2023) Altered serum bile acid profile in fibromyalgia is associated with specific gut microbiome changes and symptom severity. Pain 164:e66–e76.

Minerbi A, Gonzalez E, Brereton NJB, Anjarkouchian A, Dewar K, Fitzcharles M-A, Chevalier S, Shir Y (2019) Altered microbiome composition in individuals with fibromyalgia. Pain 160:2589–2602.

Mortensen PB, Clausen MR (1996) Short-chain fatty acids in the human colon: relation to gastrointestinal health and disease. Scand J Gastroenterol Suppl 216:132–148.

Nhu NT, Chen DY-T, Yang Y-CSH, Lo Y-C, Kang J-H (2023) Associations Between Brain-Gut Axis and Psychological Distress in Fibromyalgia: A Microbiota and Magnetic Resonance Imaging Study. J Pain Available at: 10.1016/j.jpain.2023.10.015.

Ozgocmen S, Ardicoglu O (2000) Lipid profile in patients with fibromyalgia and myofascial pain syndromes. Yonsei Med J 41:541–545.

Palomo-López P, Becerro-de-Bengoa-Vallejo R, Elena-Losa-Iglesias M, López-López D, Rodríguez-Sanz D, Cáceres-León M, Calvo-Lobo C (2019) Relationship of Depression Scores and Ranges in Women Who Suffer From Fibromyalgia by Age Distribution: A Case-Control Study. Worldviews Evid Based Nurs 16:211–220.

Ramírez-Tejero JA, Martínez-Lara E, Rus A, Camacho MV, Del Moral ML, Siles E (2018) Insight into the biological pathways underlying fibromyalgia by a proteomic approach. J Proteomics 186:47–55.

Rasouli-Saravani A, Jahankhani K, Moradi S, Gorgani M, Shafaghat Z, Mirsanei Z, Mehmandar A, Mirzaei R (2023) Role of microbiota short-chain fatty acids in the pathogenesis of autoimmune diseases. Biomed Pharmacother 162:114620.

Ruggiero V, Mura M, Cacace E, Era B, Peri M, Sanna G, Fais A (2017) Free amino acids in fibromyalgia syndrome: relationship with clinical picture. Scand J Clin Lab Invest 77:93–97.

Rus A, López-Sánchez JA, Martínez-Martos JM, Ramírez-Expósito MJ, Molina F, Correa-Rodríguez M, Aguilar-Ferrándiz ME (2024) Predictive Ability of Serum Amino Acid Levels to Differentiate Fibromyalgia Patients from Healthy Subjects. Mol Diagn Ther 28:113–128.

Rus A, Molina F, Gassó M, Camacho MV, Peinado MÁ, del Moral ML (2016) Nitric Oxide, Inflammation, Lipid Profile, and Cortisol in Normal- and Overweight Women With Fibromyalgia. Biol Res Nurs 18:138–146.

Sánchez-Domínguez B, Bullón P, Román-Malo L, Marín-Aguilar F, Alcocer-Gómez E, Carrión AM, Sánchez-Alcazar JA, Cordero MD (2015) Oxidative stress, mitochondrial dysfunction and, inflammation common events in skin of patients with Fibromyalgia. Mitochondrion 21:69–75.

Sandström A, Ellerbrock I, Tour J, Kadetoff D, Jensen KB, Kosek E (2020) Neural correlates of conditioned pain responses in fibromyalgia subjects indicate preferential formation of new pain associations rather than extinction of irrelevant ones. Pain 161:2079– 2088.

Sarzi-Puttini P, Giorgi V, Marotto D, Atzeni F (2020) Fibromyalgia: an update on clinical characteristics, aetiopathogenesis and treatment. Nat Rev Rheumatol 16:645–660.

Schmucker DL, Ohta M, Kanai S, Sato Y, Kitani K (1990) Hepatic injury induced by bile salts: correlation between biochemical and morphological events. Hepatology 12:1216–1221.

Sheng L, Jena PK, Hu Y, Liu H-X, Nagar N, Kalanetra KM, French SW, French SW, Mills DA, Wan Y-JY (2017) Hepatic inflammation caused by dysregulated bile acid synthesis is reversible by butyrate supplementation. J Pathol 243:431–441.

Slattery SA, Niaz O, Aziz Q, Ford AC, Farmer AD (2015) Systematic review with meta-analysis: the prevalence of bile acid malabsorption in the irritable bowel syndrome with diarrhoea. Aliment Pharmacol Ther 42:3–11.

Sun N, Zhang J, Wang J, Liu Z, Wang X, Kang P, Yang C, Liu P, Zhang K (2022) Abnormal gut microbiota and bile acids in patients with first-episode major depressive disorder and correlation analysis. Psychiatry Clin Neurosci 76:321–328.

Tao Y, Zhou H, Li Z, Wu H, Wu F, Miao Z, Shi H, Huang F, Wu X (2024) TGR5 deficiency-induced anxiety and depression-like behaviors: The role of gut microbiota dysbiosis. J Affect Disord 344:219–232.

Ticho AL, Malhotra P, Dudeja PK, Gill RK, Alrefai WA (2019) Intestinal Absorption of Bile Acids in Health and Disease. Compr Physiol 10:21–56.

Tour J, Sandström A, Kadetoff D, Schalling M, Kosek E (2022) The OPRM1 gene and interactions with the 5-HT1a gene regulate conditioned pain modulation in fibromyalgia patients and healthy controls. PLoS One 17:e0277427.

Traub RJ, Tang B, Ji Y, Pandya S, Yfantis H, Sun Y (2008) A rat model of chronic postinflammatory visceral pain induced by deoxycholic acid. Gastroenterology 135:2075–2083.

Wallace DJ, Hallegua DS (2004) Fibromyalgia: the gastrointestinal link. Curr Pain Headache Rep 8:364–368.

Wang Y-D, Chen W-D, Wang M, Yu D, Forman BM, Huang W (2008) Farnesoid X receptor antagonizes nuclear factor kappaB in hepatic inflammatory response. Hepatology 48:1632–1643.

Wang Z, Jiang D, Zhang M, Teng Y, Huang Y (2023) Causal association between gut microbiota and fibromyalgia: a Mendelian randomization study. Front Microbiol 14:1305361.

Ware JE Jr, Sherbourne CD (1992) The MOS 36-item short-form health survey (SF-36). I. Conceptual framework and item selection. Med Care 30:473–483.

Warren DB, Chalmers DK, Hutchison K, Dang W, Pouton CW (2006) Molecular dynamics simulations of spontaneous bile salt aggregation. Colloids Surf A Physicochem Eng Asp 280:182–193.

Whiley L, Lawler NG, Zeng AX, Lee A, Chin S-T, Bizkarguenaga M, Bruzzone C, Embade N, Wist J, Holmes E, Millet O, Nicholson JK, Gray N (2024) Cross-Validation of Metabolic Phenotypes in SARS-CoV-2 Infected Subpopulations Using Targeted Liquid Chromatography-Mass Spectrometry (LC-MS). J Proteome Res 23:1313– 1327.

Wolfe F, Clauw DJ, Fitzcharles M-A, Goldenberg DL, Häuser W, Katz RS, Mease P, Russell AS, Russell IJ, Winfield JB (2011) Fibromyalgia criteria and severity scales for clinical and epidemiological studies: a modification of the ACR Preliminary Diagnostic Criteria for Fibromyalgia. J Rheumatol 38:1113–1122.

Wolfe F, Smythe HA, Yunus MB, Bennett RM, Bombardier C, Goldenberg DL, Tugwell P, Campbell SM, Abeles M, Clark P (1990) The American College of Rheumatology 1990 Criteria for the Classification of Fibromyalgia. Report of the Multicenter Criteria Committee. Arthritis Rheum 33:160–172.

Yoneno K, Hisamatsu T, Shimamura K, Kamada N, Ichikawa R, Kitazume MT, Mori M, Uo M, Namikawa Y, Matsuoka K, Sato T, Koganei K, Sugita A, Kanai T, Hibi T (2013) TGR5 signalling inhibits the production of pro-inflammatory cytokines by in vitro differentiated inflammatory and intestinal macrophages in Crohn’s disease. Immunology 139:19–29.

Yu Y, Villalobos-Hernandez EC, Pradhananga S, Baker CC, Keating C, Grundy D, Lomax AE, Reed DE (2019) Deoxycholic acid activates colonic afferent nerves via 5-HT3 receptor-dependent and -independent mechanisms. Am J Physiol Gastrointest Liver Physiol 317:G275–G284.

Zhong S, Liu F, Giniatullin R, Jolkkonen J, Li Y, Zhou Z, Lin X, Liu C, Zhang X, Liu Z, Lv C, Guo Q, Zhao C (2023) Blockade of CCR5 suppresses paclitaxel-induced peripheral neuropathic pain caused by increased deoxycholic acid. Cell Rep 42:113386.

Zhou F, Wang X, Han B, Tang X, Liu R, Ji Q, Zhou Z, Zhang L (2021) Short-chain fatty acids contribute to neuropathic pain via regulating microglia activation and polarization. Mol Pain 17:1744806921996520.

Zhou Q, Zhang B, Verne GN (2009) Intestinal membrane permeability and hypersensitivity in the irritable bowel syndrome. Pain 146:41–46.

Zigmond AS, Snaith RP (1983) The hospital anxiety and depression scale. Acta Psychiatr Scand 67:361–370.

