## Supplementary material for "Microbially Produced Bile Acids are Associated with High Levels of IgG Autoantibodies and Worse Mental Well-being in Fibromyalgia Subjects"

### Supplementary figures

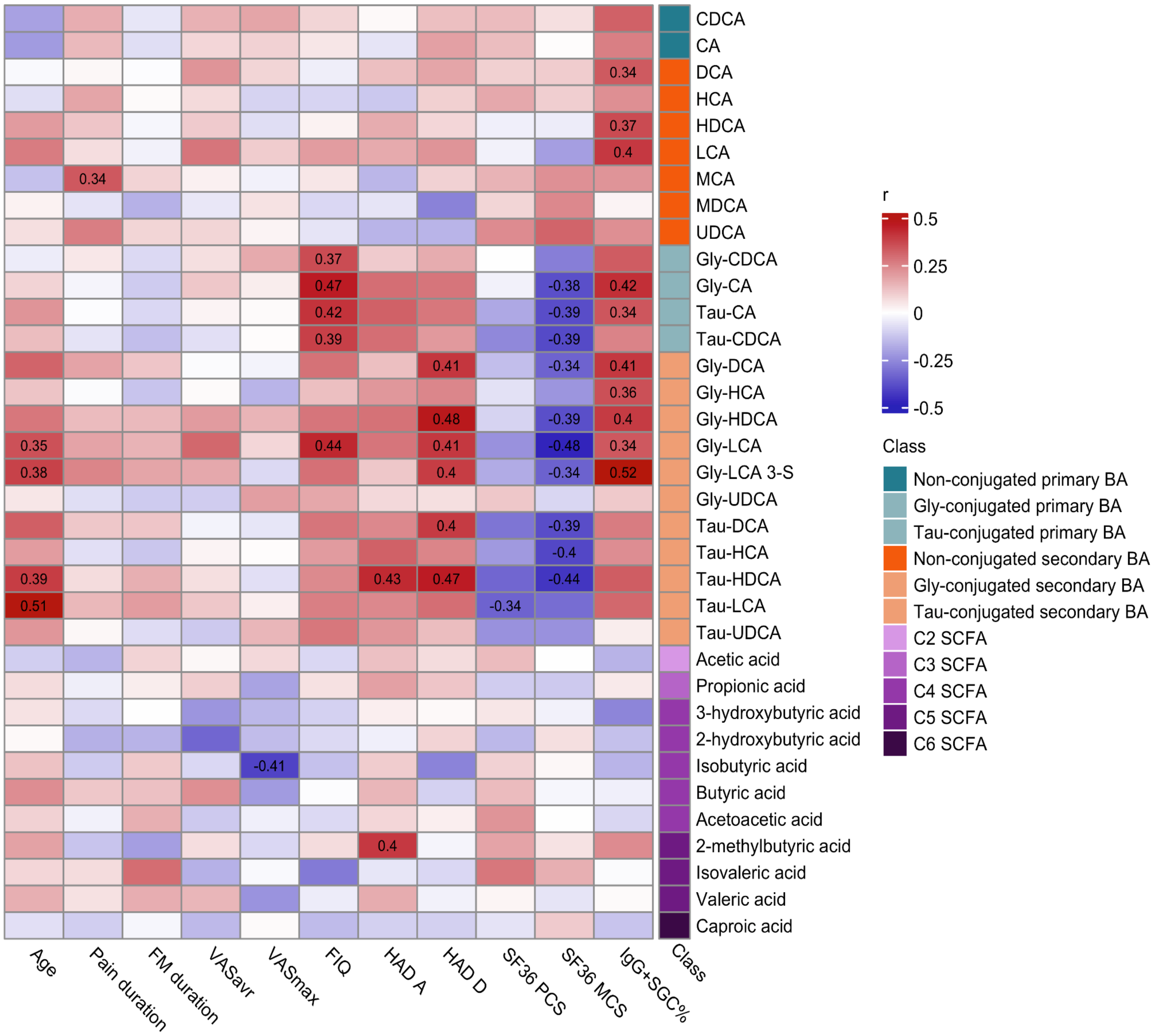


**Supplementary Figure S1. Fibromyalgia (FM) subjects have associations between conjugated bile acids (BAs) and mental well-being (SF36 MCS).** FM subjects' BA and short-chain fatty acid (SCFA) concentrations were correlated to demographic and clinical data. Secondary BAs DCA, LCA, and HDCA, as well as glycine (gly)-conjugated DCA, HDCA, HCA, CA, LCA 3-S, and LCA, were positively correlated to anti-SGC IgG levels (IgG+SGC%). Gly-CA, Tau-CDCA, Tau-CA, Gly-DCA, Gly-HDCA, Gly-LCA, Gly-LCA 3-S, Tau-DCA, Tau-HCA, and Tau-HDCA were associated with worse mental well-being (summary of the mental component of the short form 36 health survey questionnaire; SF36 MCS). Gly-conjugated DCA, HDCA, LCA 3-S, and LCA and taurine (tau)-conjugated DCA and HDCA concentrations were positively correlated to depression scores (Hospital anxiety and depression scale; HAD-D). The correlation analysis was performed with Spearman's rank or Pearson, where applicable, correlation. A positive correlation is colored in red, and a negative in blue. The numbers shown are the correlation coefficients for significant ($P < 0.05$) correlations. The class legend illustrates what type of BA it is (primary, secondary, and conjugated or not) or the number of carbons (C2 - C6) the SCFA has.
